# The damage-independent evolution of ageing by selective destruction

**DOI:** 10.1101/2022.05.03.490465

**Authors:** James Wordsworth, Hannah O’ Keefe, Peter Clark, Daryl Shanley

## Abstract

Ageing is currently believed to reflect the accumulation of molecular damage due to energetic costs of maintenance, as proposed in disposable soma theory (DST). Here we use agent-based modelling to describe an alternative theory by which ageing could undergo positive selection independent of energetic costs. We suggest that the selective advantage of aberrant cells with fast growth might necessitate a mechanism of counterselection we name selective destruction that specifically removes the faster cells from tissues, preventing the morbidity and mortality risks they pose. The resulting survival advantage of slower mutants could switch the direction of selection, allowing them to outcompete both fast mutants and wildtype cells, causing them to spread and induce ageing in the form of a metabolic slowdown.

Selective destruction could therefore provide a proximal cause of ageing that is both consistent with the gene expression hallmarks of ageing, and independent of accumulating damage. Furthermore, negligible senescence would acquire a new meaning of increased basal mortality.

## Introduction

Ageing is a multifactorial process (Kirkwood 2005; López-Otín et al. 2013). However, nearly all existing theories ultimately require molecular damage to be the initial cause (Kirkwood 2005). In disposable soma theory (DST) for example, Kirkwood (1977) posited that investing in maintenance is energetically costly, and therefore molecular damage could be allowed to accumulate if that energy could be better utilised in other processes with greater impact on fitness (e.g. growth and reproduction). While Kirkwood (2005) suggested multiple molecular mechanisms were likely to be involved in ageing, including telomere attrition, protein aggregation, and mitochondrial dysfunction, all the suggested mechanisms were based on molecular damage, and ageing would thus reflect “multiple kinds of damage “. However, if that is true then at its core, ageing reflects only one single cause: that organisms do not invest sufficiently in maintenance to prevent damage accumulation. Indeed, few hypotheses suggest how ageing could result without damage accumulation.

The main exceptions are mutation accumulation theory (MAT) and antagonistic pleiotropy (AP). The former suggests that high extrinsic mortality in the wild would allow for ageing-inducing mutations to persist due to the resultant declining force of selection with age (Medawar 1952). Initially, the idea was attractive because ageing was not believed to occur in the wild (Medawar 1952). However, it is now accepted that ageing does occur in the wild in multiple species (Nussey et al. 2013). In contrast to MAT, AP suggests that pro-ageing mutations offer a fitness advantage earlier in life when selection is stronger (Williams 1957). As with MAT, there are also multiple single nucleotide polymorphisms (SNPs) in humans that may contribute to early life fitness at the cost of fitness later in life (Rodríguez et al. 2017), and predictably, these mutations have detrimental effects significantly earlier. However, while such SNPs may explain the onset of some diseases in some individuals, it is more difficult to identify ancient mutations that are now universal, inducing ageing via a consistent mechanism across species and organisms. As such, these theories currently provide a clearer view of the evolutionary framework that may allow ageing to evolve rather than a comprehensive biological mechanism.

Here, we outline an antagonistically pleiotropic biological process that could induce ageing independently of damage accumulation and the energetic costs of maintenance. As such, we believe selective destruction theory (SDT) provides the first comprehensive mechanism providing ageing organisms with a fitness advantage over similar organisms undergoing negligible senescence without dependence on damage accumulation or energetic costs. However, it is not mutually exclusive with damage-centric theories.

### Theory outline

It is only recently that the plethora of selective forces acting within tissues are becoming appreciated. Just as cancer cells undergo positive selection due to their fast growth and limited response to control, allowing them to spread and ultimately cause death, non-tumour cells can also undergo permanent or semi-permanent changes that give them a selective advantage and allow them to spread at the expense of non-mutant (wildtype) cells (reviewed by Kakiuchi and Ogawa (2021)). This *within-tissue* selection could therefore affect the fitness of organisms, and influence natural selection. Indeed, an important modelling study by Nelson and Masel (2017) even suggested that such forces might make ageing inevitable.

Here we are interested in changes affecting growth and proliferation. These could reflect mutations or epigenetic changes, but the altered cells will henceforth be referred to as mutants for simplicity. Such mutants are either aberrantly sensitive (AS) or aberrantly resistant (AR) to growth signals compared with wildtype cells. While both AR and AS mutants reduce tissue functionality by responding incorrectly to environmental cues, AR mutants will grow and proliferate slower than wildtype cells, putting them at a selective disadvantage, whereas AS mutants will grow and proliferate faster, giving them a selective advantage over wildtype cells and AR mutants. AS mutants are therefore a threat to tissue homeostasis in a way that AR mutants are not. If they are not controlled or removed, they will outcompete the wildtype cells and may become the dominant cell type (Figure 1A).

**Figure 1.**
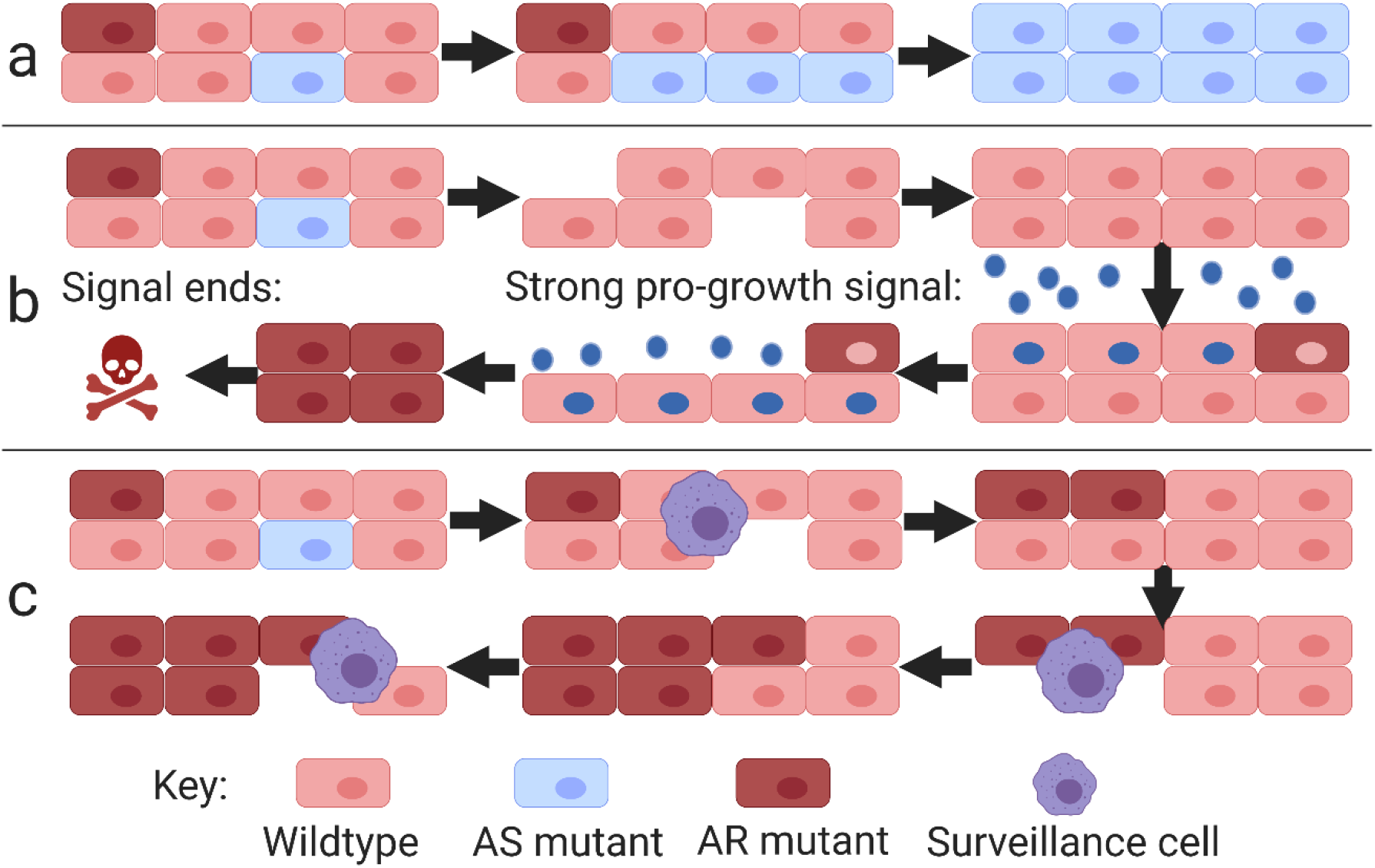
Tissue development with different types of mutant control. (A) shows no mutant control, the expansion of AS mutants and their takeover of the tissue. (B) shows autonomous control. Each cell recognises its own sensitivity and can kill itself when its stimulus suggests that it has undergone mutation and is no longer responding correctly to the environment. As shown on the top half, such mechanisms efficiently remove both types of mutant in a constant environment, but if the environment changes as shown in the bottom half, inducing a strong pro-growth stimulus for example, the cells are unaware that their neighbours have had a similar response and assume the changes reflect their own aberration. They therefore die, leaving only the mutants which the aberrant stimulus makes appear wildtype. If the stimulus recedes, these cells too will die as they now correctly identify themselves as mutant again. (C) Cells compare themselves to their neighbours, preferentially removing faster cells. This allows the AR mutants to slowly accumulate (top) until there are areas in the tissue where their dominance causes removal of wildtype cells (bottom). AR, aberrantly resistant; AS, aberrantly sensitive.

Even beyond their selective advantage, AS cells represent an increased threat to tissue homeostasis. The accelerated growth and division of AS cells would be associated with a faster mutation rate, as can be seen under extreme examples such as RAS and Rapidly Accelerated Fibrosarcoma (RAF) mutations which induce a state of hyperproliferation associated with high levels of DNA damage (Janda et al. 2002). While these extreme mutations induce senescence (Di Micco et al. 2006), for reasons described below, mutations with milder effects on growth cannot autonomously stop their own growth. Therefore, if left uncontrolled, such mutants could undergo successive mutation and transformation, increasing the risk of tumorigenesis.

Another potential risk is fibrosis, which is a highly metabolic process requiring activation of growth pathways such as mTORC1 signalling to both induce rapid cell division and produce the large amounts of extracellular matrix (ECM) proteins (Woodcock et al. 2019). In some mesenchymal cell types, AS mutants with faster metabolism and more active growth pathways could therefore be expected to increase the risk of fibrosis. For example, insulin-like growth factor-1 (IGF-1) is a key growth factor inducing growth and proliferation, which also stimulates the survival and activation of fibroblasts, causing differentiation to highly fibrotic myofibroblasts (Choi et al. 2009; Hung et al. 2013). AS mutants with more active IGF-1 signalling would therefore proliferate and activate in response to weaker signals, making it more likely that they would reach the critical mass required for fibrosis (Adler et al. 2020). Patients with systemic sclerosis and idiopathic pulmonary fibrosis (IPF) both have high levels of serum IGF-1 (Hamaguchi et al. 2008), indicating that these lethal diseases involve increased activation of pathways associated with AS mutants. Notably, around 45% of all deaths in the developed world are ascribed to chronic fibroproliferative diseases (Wynn 2007).

There are also many other examples where cell function might be upregulated in AS mutants as many cell types link their specific functional output (such as hormones or metabolites) to growth pathways. This reflects that growth and proliferation are usually part of the mechanism to increase long-term production/activity so pathways which increase one will increase the other (Karin and Alon 2017). For example, β cells proliferate and produce insulin when blood glucose is high, and trigger apoptosis and inhibit insulin production when glucose is low. The insulin production pathway is therefore intertwined with the growth and proliferation pathways (Rhodes and White 2002) so both can be up or downregulated together in response to stimulus. However, as the human body requires blood glucose to remain at around roughly 5 mM, Karin and Alon (2017) identified that mutants sensing blood glucose at an incorrectly high level would quickly cause death if they were allowed to spread. This would happen for two reasons. Firstly, these mutants hypersecreting insulin can lower blood glucose below the point at which the rest of the body’s tissues can sustain respiration. Secondly, the wildtype cells sensing the correct level of glucose would undergo apoptosis to reduce insulin production, hastening the spread of mutants and the lethal drop in blood glucose. Thus, any cell type that utilises cell growth and division as mechanisms to regulate functional output, and whose functional output relies on keeping metabolites, proteins, and even other cell types within a certain range, is threatened by AS mutants. Even if the outcomes are not lethal, as they are for fibrosis, insulin production, and other pathways such as calcium homeostasis, they will not promote good health or bodily function.

AS mutants will therefore promote morbidity and mortality **through cancer, fibrosis, and overactivity (CFOA)**. From this, we conclude that organisms would gain a fitness advantage from controlling AS mutants through maintaining better tissue homeostasis. The stricter the control mechanisms, the longer an organism could hope to retain wildtype functionality and avoid CFOA. By contrast, AR mutants provide little threat to tissue function. Their slow growth will make them naturally outcompeted, but even if they accumulated, their reduced metabolic function would not induce the same threat as AS mutants because they would produce less of their own specific output, merely requiring wildtype cells to work harder to maintain tissue function. We suggest that this asymmetry between the threat posed by AS and AR mutants might necessitate a control mechanism tailored toward removing AS cells and the immediate threat they pose.

There are two possible mechanisms of controlling mutants with a selective advantage, both requiring a marker of phenotypic change: as here we are concerned with sensitivity to growth, we have termed this a **growth marker (GM)**.

Firstly, cells could autonomously recognise that the absolute concentration of the GM they are expressing is too high and induce their own apoptosis. Oncogenic mutations in RAS, AKT, PI3K, and multiple other mitogenic molecules have been shown to induce senescence and apoptosis depending on cell type and severity (Liu et al. 2018). Hypersecreting β cell mutants also undergo apoptosis (termed glucotoxicity) (Karin and Alon 2017).

Importantly, autonomous mechanisms allow both AS and AR mutants to be removed with equal efficiency, as cells recognise their own absolute value of GM (Figure 1B). However, such mechanisms are intrinsically dangerous: in the case of β cells, glucotoxicity in hunter gatherers would have removed only dangerous mutant cells producing too much insulin, but modern diets rich in calories and refined sugars are inducing glucotoxicity in wildtype cells which are producing high levels of insulin to remove the high levels of glucose in the blood, thus resulting in pancreatic destruction and type II diabetes (Karin and Alon 2017). The danger arises because **autonomous mechanisms are incapable of distinguishing between aberrant cells and aberrant conditions**, so any autonomous mechanism must only activate in conditions that are highly unlikely to occur in nature. Consistently, type II diabetes was likely a rare condition before the introduction of refined sugars (Johnson et al. 2017).

Secondly, mutants could be controlled via a non-autonomous method via comparison with the surrounding tissue. For example, tumour cells are distinguished from surrounding tissue by natural killer (NK) cell recognition of MHC class I chain-related protein A (MICA) levels (Møller et al. 2020). In the case of β cells, only the most extreme hypersecreting mutants induce glucotoxicity, so there is still a range of mutants which cannot be removed this way, but still have a selective advantage over wildtype cells. Korem Kohanim et al. (2020) suggested a system of immune control called the autoimmune surveillance of hypersecreting mutants (ASHM): autoimmune T cells would compare the level of pro-insulin expressed by cells before killing the cell expressing the highest levels. Thus, if a mutation increasing insulin production has occurred in a cell, then it would likely be killed first.

For AS mutants, when correctly calibrated, an ASHM-like mechanism could correctly discern aberrant cells from aberrant conditions, allowing even low level mutants to be removed this way,and thus returning the selective advantage to wildtype cells, as shown in Figure 1C (top). However, as also shown in Figure 1C (bottom), such **selective destruction of the fastest growing cells** is incapable of removing the slow growing AR mutants, which will slowly replicate over time until they begin to outnumber wildtype cells in certain areas. At this point, the immune cells would then recognise the AR mutants as wildtype and the wildtype as AS mutants, providing a selective advantage to the AR mutants. Their resultant accumulation could cause tissue robustness to decline as the average functionality of cells slows, which we propose could cause ageing. Importantly, in the model of hypersecreting cells described by Korem Kohanim et al. (2020), the spread of hypo-secreting mutants was unlikely because such cells preferentially induced apoptosis aspart of the mechanism to raise blood glucose (Topp et al. 2000). Severe AR mutants may also undergo apoptosis due to lack of mitogenic signals, but moderate mutants are likely to persist if the soma uses a system of selective destruction for mutant control. Attempting to remove AR mutants by either autonomous or comparative mechanisms could have serious repercussions: ASHM, for example, has been implicated in type I diabetes through the immune destruction of healthy β cells (Korem Kohanim et al. 2020) because chance events such as infection lead to the identification of healthy β cells as hypersecreting mutants, causing the destruction of the pancreas.

A common problem for antagonistically pleiotropic mechanisms is that evolution can separate the benefits from the costs. However, we hypothesised that no changes to the process of selective destruction could result in a more favourable outcome for the organism. The options would be threefold:

1. Spread of AS mutants resulting from their selective advantage. The result is death from CFOA at an early age.
2. Autonomous control of AR and AS mutants even for low level changes. The result is death from *en masse* apoptosis and senescence when changes in environmental conditions cause wildtype cells to be mistaken for mutants.
3. The **selective destruction (SD)** of AS cells which allows AR mutants to spread. The result is slow functional decline (ageing).

From the three possible outcomes, ageing would provide the greatest fitness advantage, while the rate of ageing would depend on the severity of the SD. To address whether this was true in practise we constructed a series of models to address if:

1. SD would prove better at mutant control than **unselective destruction (UD)** i.e. equal attempt to remove both AS and AR mutants (model 1).
2. If implementing SD would induce ageing as predicted via the spread of AR mutants (model 1).
3. Under what conditions ageing by SD would undergo positive selection (model 2).

If SD involved an immune surveillance mechanism similar to that described by Korem Kohanim et al. (2020) for the comparative control of hypersecreting mutants, it would likely work mainly via inducing apoptosis. However, we considered that while such a process might work well for the regulation of the small populations of cells in pancreatic islets, it was unlikely to be the central mechanism across multiple organs and tissues maintained by selective destruction, particularly among simpler creatures with more primitive immune systems that nonetheless still age.

Instead, we hypothesised that selective destruction could be implemented by juxtacrine and paracrine signals from cells within the same tissue. Speculatively, neighbouring cells could communicate their relative level of metabolism by the levels of GM, and influence each other’s fate accordingly, with slower cells suppressing the growth potential of faster cells through epigenetic modification, senescence, or apoptosis.

Notch signalling controls juxtacrine communication, and consistently, Notch mutations are associated with the clonal expansion of leukaemia cells (Kimura et al. 2019) as well as skin and lung squamous cell carcinomas (Wang et al. 2011), which may implicate that these cells are escaping attempts of their neighbours to suppress their clonal advantage, highly consistent with SDT. Indeed, evidence suggests that Notch mutations aid tumour formation via non-autonomous signals from the tumour microenvironment (Demehri et al. 2009), while Notch signalling plays a key role in senescence by mediating juxtacrine signalling (Hoare et al. 2016). Thus, in this theoretical study, we attempt to demonstrate the plausibility of ageing by juxtacrine SD.

## Materials and Methods

### Model 1: Tissue Selection

NetLogo (version 6.2.0) (Wilensky 1999) was used to construct a series of agent-based models on a two-dimensional grid of cells measuring 33 by 33 for a total of 1089 cells of a single cell type. At the beginning, all cells were considered to have wildtype sensitivity to growth and proliferation signals. Then over a series of cycles, the cells could mutate to become either AS or AR mutants as measured on a scale of 0-4: AR mutants had sensitivity values of 0 and 1, the wildtype a value of 2, and AS mutants values of 3 and 4, with 0 being the least sensitive and 4 being the most sensitive. This is considered to represent the viable range of mutations: cells with lower sensitivity would not receive sufficient mitogenic signals to survive, while cells with higher sensitivity could be removed by autonomous mechanisms (as they are sufficiently aberrant that they are unlikely to reflect any natural conditions), as shown in Figure 2A. Importantly, the size of this range and the relative advantage of cells with higher sensitivity can be varied using the *Fill* variable, as described below.

**Figure 2.**
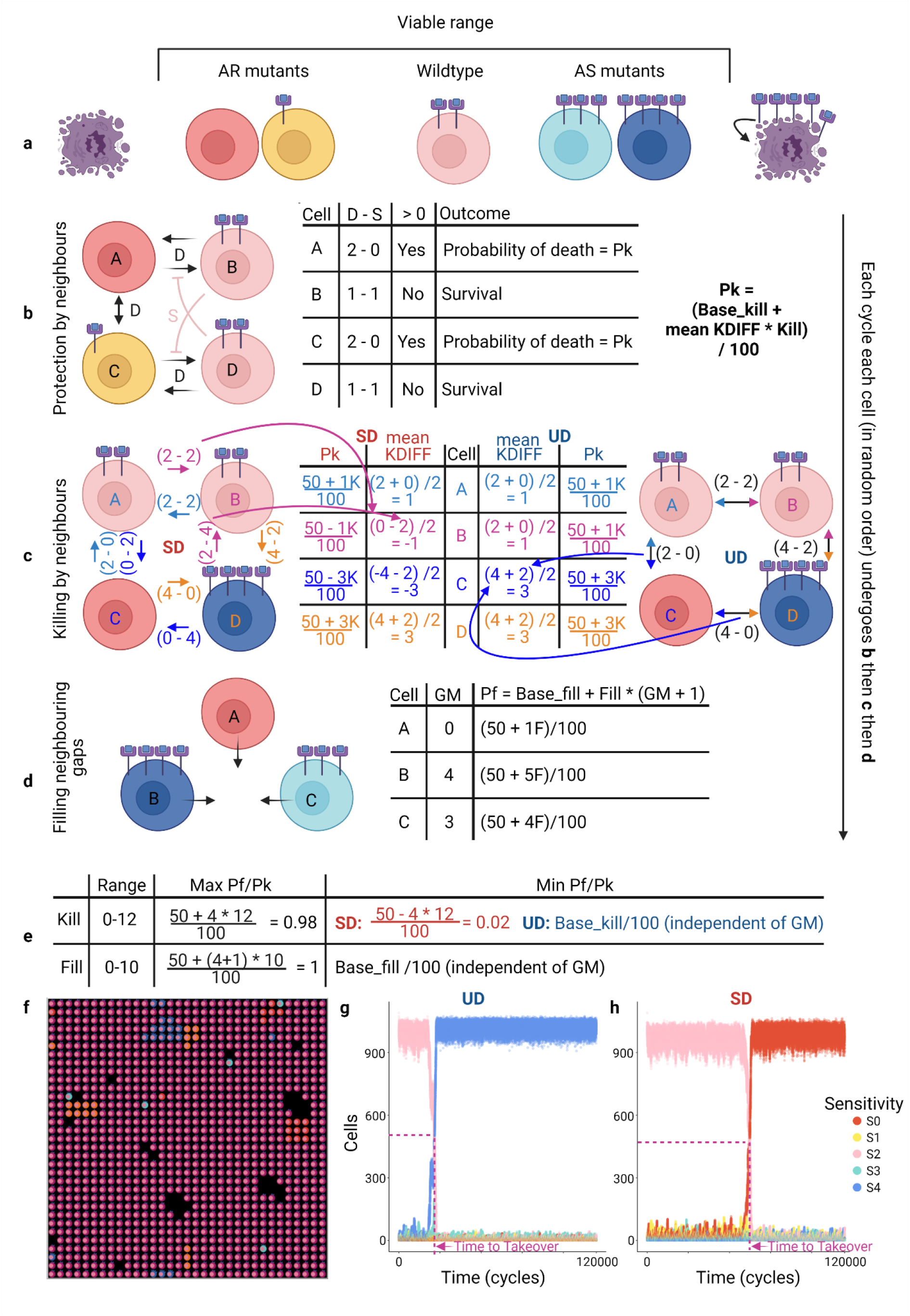
Tissue Selection Model Diagram (A) Schematic of hypothetical viable range of mutations conferring different sensitivity to growth and proliferation stimuli, with changes outside this range inducing death. Cell colours and number of receptors indicate the relative production of growth marker (GM), which reflects their sensitivity to growth stimuli (see E for more details). (B) Each cycle, each cell counts the number of neighbours with the same GM (S in diagram) and those with different GM (D in diagram). If a cell has more different than similar neighbours (D – S > 0), then it has a probability, Pk of being killed that cycle, but if D – S ≤ 0 it receives sufficient survival signals to survive that cycle. The diagram shows this calculation for example cells with two neighbours. (C) The probability that a cell (with D – S > 0) will be killed differs between the two models. The diagram shows the calculation made by a cell with two neighbours. In SD, each cell minuses both neighbours’ GM from its own. As shown in the table, the average of these two values forms the KDIFF which is multiplied by the variable Kill (K in the table) and added to the base_kill variable (50 in the table). The probability that the cell will die, Pk, is the result/100. Thus, in SD cells surrounded by neighbours with more GM have negative KDIFF values, which decreases their likelihood of death. Conversely, in UD (right) it is only the magnitude of the difference in GM that counts, so KDIFF is always positive, and the Pk for slow cells with fast neighbours is the same as vice versa. (D) If the cell survives, then it has probability Pf of filling any neighbouring gaps, which is determined by (Base_fill [50 in table] + Fill *(GM of the cell + 1))/100. Thus, cells with greater sensitivity are more likely to fill gaps, and the level of advantage gained is determined by the variable Fill. (E) The range of Fill and Kill values used in our simulations created a full spectrum of Pf and Pk values respectively, reflecting tissues that range from completely insensitive to GM (i.e. no mutation affects growth rate) to tissues where the strongest mutations gained the maximal possible advantage. The Fill values determine the autonomous control of mutants, with higher Fill values reflecting lower autonomous control which allows the survival of mutants with greater changes in sensitivity. The Kill values determine non-autonomous control, with higher Kill values producing greater probability that cells are killed by their neighbours (and reduced probability for slower cells in SD). (F) Example run of SD tissue model. Magenta cells are wildtype S2, Cyan and Blue are AS mutants with S3 and S4, respectively. Orange and red cells are AR mutants with S1 and S0, respectively. Kill 2 Fill 2 cycle 1020. (G and H) show the abundance of cells with different GM in example runs for UD and SD respectively. Time to takeover is demonstrated for each. (G) UD Fill4 Kill4 replicate 10. (H) SD Fill 2 Kill 7 replicate 7. AR, aberrantly resistant; AS, aberrantly sensitive; D, Different (cells with different sensitivity); F, Fill (variable); K, Kill (variable); S, Same (Cells with same sensitivity); SD, selective destruction; UD, unselective destruction.

Gaps were created in part by juxtacrine killing (see below), but also by cell death from independent factors determined by probability of ~0.011 per cell per cycle, with ~0.006 probability that surrounding cells would also die. To accelerate the model behaviour, mutation had a probability of 0.01 per cell per cycle, with additional chance of mutating during division of 0.02, which could happen before or after division (affecting one or both daughter cells). Each cycle, cells were run in random order. First, cells compared their GM to their neighbours. Although such a phenomenon is not well documented in the literature, we considered that such juxtacrine regulation of survival would be most efficient (causing the least unnecessary or detrimental cell death) if it advantaged the wildtype cells as much as possible. Therefore, while our model assumes that cells cannot intrinsically know what is the ‘correct’ (or wildtype) level of GM, we considered that the initial numerousness of wildtype cells compared to mutants could be used to prevent their deaths if cells with similar GM provided a survival signal, exemplified in Figure 2B. Therefore, each cycle cells would compare the number of neighbouring cells with the same level of GM as themselves to those with different levels, and only if the latter outweighed the former might the cell be killed that cycle. Without this 0.01 protection, it would be considerably harder to prevent the spread of either AS or AR mutants. Therefore, only if a cell was not sufficiently similar to its neighbours, the probability of being killed, Pk, was then determined (equation 1).

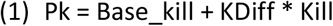

The *Base_kill* determined the probability of cells being killed by their neighbours independent of their GM. The *Kill* variable determined the magnitude of the change from *Base_kill* that would be applied to cells according to the *KDiff* value, which reflected the relative GM of a cell compared to its neighbours (equation 2).

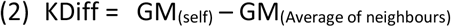

We considered there were several reasons that greater difference might result in higher probability of killing: smaller changes were more likely to be obscured by environmental differences, and larger changes could reflect increasingly aberrant and therefore dangerous cells.

We then used either *KDiff* or |*KDiff*| to compare the two different mechanisms of mutant control: SD, which affords protection to slower cells, used *KDiff*, so that if the cell had less GM than its neighbours *KDiff* would be negative (subtracted from *Base_kill*, reducing *Pk*), while UD utilised

|*KDiff*|, killing slower and faster cells with equal proficiency, as exemplified in Figure 2C. Importantly, the only difference between SD and UD was that SD made it more difficult to kill slower cells, while both models removed faster cells with equal proficiency. *KDiff* was then affected by a multiplier variable *Kill*.

If cells survived killing by their neighbours, they could fill neighbouring gaps according to probability,

*Pf*, that reflected the relative sensitivity to growth (equation 3).

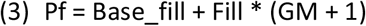

The *Base_fill* determined the probability of a cell filling a gap independent of sensitivity to growth (as indicated by GM). The additional probability was then determined by sensitivity (GM) *multiplier variable, *Fill*, which thus determined the magnitude of the growth advantage accorded to cells with different GM (we used GM +1 so GM=0 did not negate the *Fill* value), as exemplified in Figure 2D.

The multiplier variables *Fill* and *Kill* determine the level of advantage given to cells with different sensitivity. Higher *Fill* values enlarge the difference between cells with high growth sensitivity and those with low, in effect increasing the viable range of mutations (before autonomous killing can remove faster cells). Higher *Kill* values have different effects in the two different models. In UD, they will result in higher probability that cells with neighbours of different sensitivity will be killed by them (increasing *Pk*). The same is true in SD except if the cell is slower than its neighbours, in which case higher *Kill* values will reduce *Pk* (see Figure 2C). The range of *Fill* and *Kill* values used in our simulations create a large spectrum of *Pf* and *Pk* probabilities, see Figure 2E, reflecting theoretical tissues that range from completely insensitive to GM (i.e. no mutation affects growth rate) to tissues where the strongest mutations gained the maximal possible advantage. Thus, the highest *Fill* values would represent organisms with no autonomous control of mutants, and the lowest values would reflect complete autonomous control of mutants, with values in between corresponding to every possible viable range of growth mutations. Similarly, the lowest *Kill* values result in juxtacrine killing that is independent of GM, and the highest values result in almost certain killing for cells with the greatest difference in sensitivity to their neighbours (or almost certain survival for slower cells in SD). It is near inconceivable that any lifeform would lie outside this range.

Models for SD and UD were run for 120,000 cycles each with twelve replicates (example in Figure 2F). We then calculated time to takeover as the number of cycles before any type of mutant first outnumbered wildtype cells and subtracted this from the total run time so that values centred around zero. We called this speed of takeover. As takeover only ever went in a single direction (AS or AR), this accurately reflected the level of advantage for the variable values used. Example runs are shown in Figure 2G and H.

### Model 2: Gene Selection

We constructed a NetLogo model of an agent population with three genes for mutant control, each with two alleles that either strengthened SD (S alleles) or relied on UD (U alleles). Having no SD alleles would result in AS dominance and increasing risk of CFOA as AS cells spread, one S allele would result in wildtype dominance producing negligible senescence, and two or more would result in AR mutant dominance and ageing. Three S alleles would induce faster ageing than two, but each S allele (1-3) would reduce the risk of CFOA by accelerating the clearance of AS mutants, as summarised in Figure 3.

**Figure 3.**
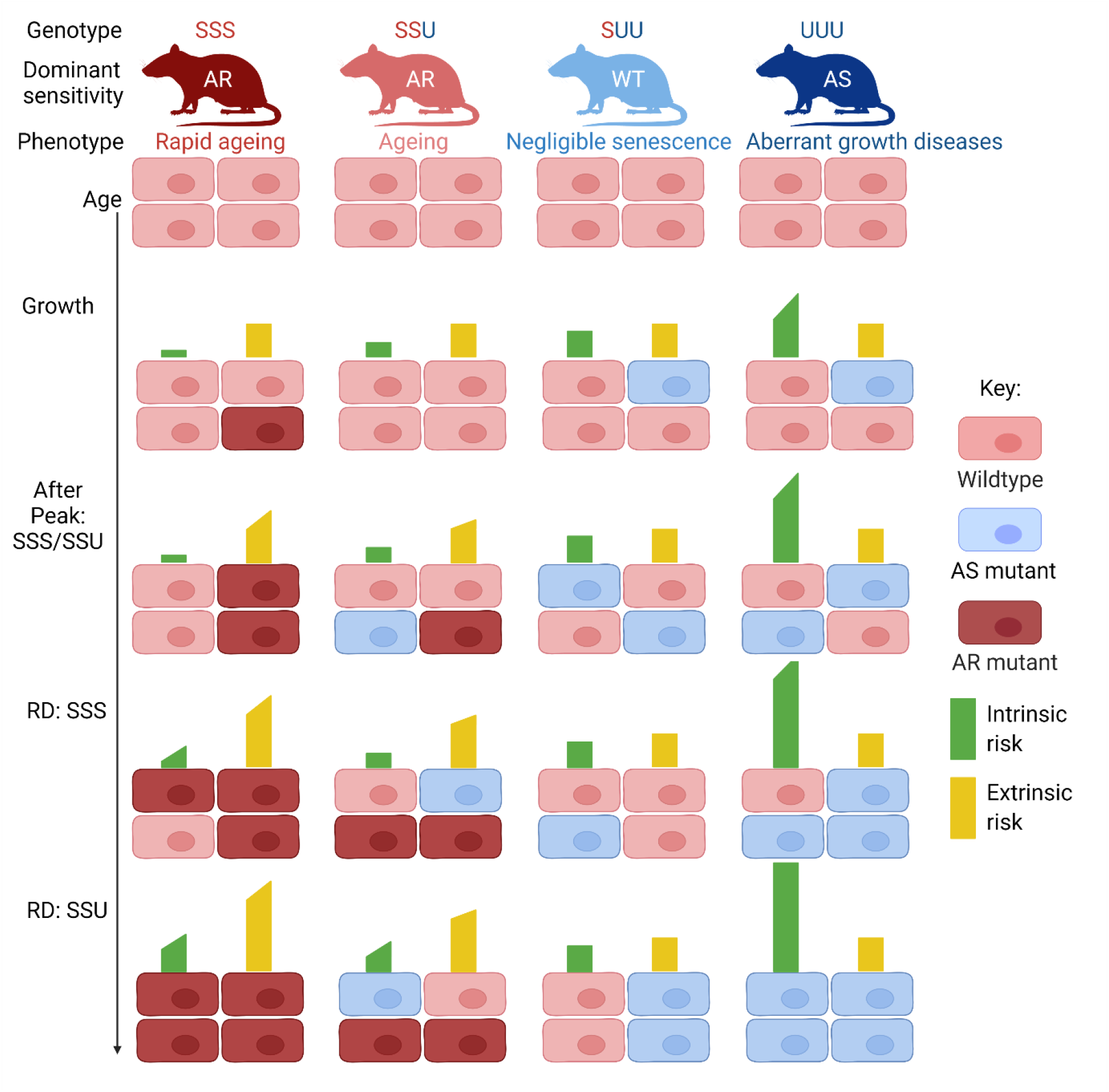
Relative risks for different allelic combinations over time. The genotype reflects the combinations of two alleles, S for SD and U for UD, for three genes. More S alleles strengthens SD, shifting selection to counterselection and changing the dominance from the naturally dominant AS cells through wildtype and finally toward AR dominance. The phenotypes of the resultant mutant spread are shown. The cells, which are not included in this model, represent the effects on tissue homeostasis, as shown by the relative levels of mutant cells. As time passes, the genotypes (and the resultant dominant sensitivity) have different effects on intrinsic and extrinsic risk, which are shown by the bars above the cells, first during the period of initial growth, then after the peak age (variable Peak) where ageing organisms begin to undergo physical decline, and finally in late life where ageing organisms have increased intrinsic risk reflecting age-related disease (RD, Rapid decline). A flat bar indicates the risk is not changing at that age, whereas a sloping bar indicates risk is increasing linearly over time. The heights of the bars are comparable between bars of the same colour, as is the steepness of the slope (for increasing rates). AR, aberrantly resistant; AS, aberrantly sensitive; RD, Rapid decline; WT, wildtype.

Each cycle agents moved a random distance (1-4) at a random heading. If two agents of sexual maturity, as determined by the variable *Mating age*, occupied the same space, then they produced an offspring, which gained a mixture of the six alleles for the three genes from the two parents, with a 1 in 400 chance of an allele mutating (from S to U or U to S). All runs were 100,000 cycles in length and had 12 replicates. The starting population size was 50, and to maintain population stability, extrinsic mortality increased with population size.

In defining intrinsic and extrinsic mortality, we were aware of the continuing debate about the relevance of extrinsic mortality to ageing (de Vries et al. 2022; Moorad et al. 2019). However, there is general consensus, even if age-independent extrinsic risk cannot impact selection for senescence, that condition/age-dependent extrinsic risk could (Moorad et al. 2019). For our purposes, age-dependent extrinsic risk would reflect the impact of intrinsic changes such as ageing on the probability of succumbing to extrinsic risks, and will be referred to as extrinsic risk. To keep the model as simple as possible, we assumed that CFOA would have little effect on extrinsic risk, with all three components representing acute intrinsic attacks that affect mortality independently of extrinsic threats. Ageing, by contrast, would firstly increase extrinsic risk by reducing the capacity of organisms to respond quickly and robustly to their environment. This would commence at an age determined by the variable *Peak*, then at a later age, determined by the variable *Rapid decline* (RD), ageing would also impact intrinsic risk through age-related diseases. As shown in Figure 3, all increases in risk were linear, reflecting the equations in Supplementary Table 1. The survival (function) of an individual then reflected the relative importance of intrinsic and extrinsic factors, which we termed the survivability. A survivability value of Z means an X in Z chance of death at each cycle, where X is the risk value of an individual within the population (determined by its genotype, see Figure 3). In negligibly senescing populations, X was determined by population size for extrinsic survivability, and number of S alleles for intrinsic survivability. In ageing populations, age beyond peak fitness (*Peak*) and *Rapid decline* also affected extrinsic and intrinsic X, respectively. As intrinsic and extrinsic survivability (Z values) increase, the impact of the risk (X values) is reduced, allowing us to construct a landscape of survivability values which varied the relative importance of intrinsic and extrinsic factors.

Organisms had to reach *Mating age* to reproduce and then land on the same space as another organism (also at mating age) to produce offspring, and S and U alleles could arise by mutation during reproduction.

Data were analysed using R (version 3.6.3). Regression analysis utilised the linear model fitting function lm(Mean ~ Time + LifeHistory, data=SIGTEST) from stats (version 3.6.2). Two-tailed T-tests were carried out using the pairwise_t_test function comparing each *Kill-Fill* combination between two conditions.

Calculations for risk: For tumour risk value (TRV), only mutants that persisted for at least four cycles were considered to have any risk of oncogenic transformation as cells must undergo successive mutations to become transformed. Each cycle after the fourth confers increasing risk of transformation, calculated by x + 0.1 *number of clones, where x is the previous risk coefficient (starting at 1). For fibrosis risk value (FRV), risk at each cycle was calculated as the sum of the squares of the size of the clusters. In both cases, additional weight was applied to mutants with sensitivity 4 over those with sensitivity 3 as faster mutants were considered more dangerous.

Models are publicly available on Fairdom Hub:

SD: 10.15490/fairdomhub.1.model.799.1

UD: 10.15490/fairdomhub.1.model.800.1

SD with conversion: 10.15490/fairdomhub.1.model.801.1

Gene selection model: 10.15490/fairdomhub.1.model.802.1

Data are available upon request.

## Results

### Selective destruction prevents AS mutant takeover

Running the simulations of our tissue model for up to 120,000 cycles demonstrated one of three outcomes depending on the conditions:

- Wildtype cells would retain dominance throughout the simulation
- AS mutants would spread and eventually outnumber wildtype cells
- AR mutants would spread and eventually outnumber wildtype cells

As expected, once an AS or AR mutant had become dominant it would only be superseded by a more extreme mutant (going from 3 to 4 or 1 to 0). Return to wildtype or oscillating from AS to AR never occurred. We could therefore compare the direction of takeover and the rate and at which it occurred between SD and UD. We therefore looked across the entire spectrum of *Fill* and *Kill* values (Figure 2E), with higher *Fill* values giving additional growth advantage to more sensitive cells (Figure 2D), and higher *Kill* values resulting in a greater chance that cells with different sensitivity to their neighbours are killed by them (Figure 2C). However, in SD the latter only applied if the cell was more sensitive than its neighbours, while less sensitive cells became less likely to be killed with higher values of *Kill* (Figure 2C). For each combination of *Kill* and *Fill* we used, the time to takeover (see examples Figure 2G and H) was converted to speed (see Methods) and the mean values plotted. As shown in Figure 4A, UD with even the highest values of *Kill* (resulting in high probabilities of death for all mutants) proved incapable of preventing AS mutant takeover in all but the lowest values of *Fill* (where AS mutants received the lowest increases in probability of filling gaps). AS mutants that escaped killing long enough (by chance) to proliferate into a protective niche (where their similarity to each other prevented them being killed) quickly allowed them to become dominant. Whereas SD, shown in Figure 4B, proved a significantly more effective AS mutant control mechanism at several values of *Kill* as determined by two-tailed T-tests.

**Figure 4.**
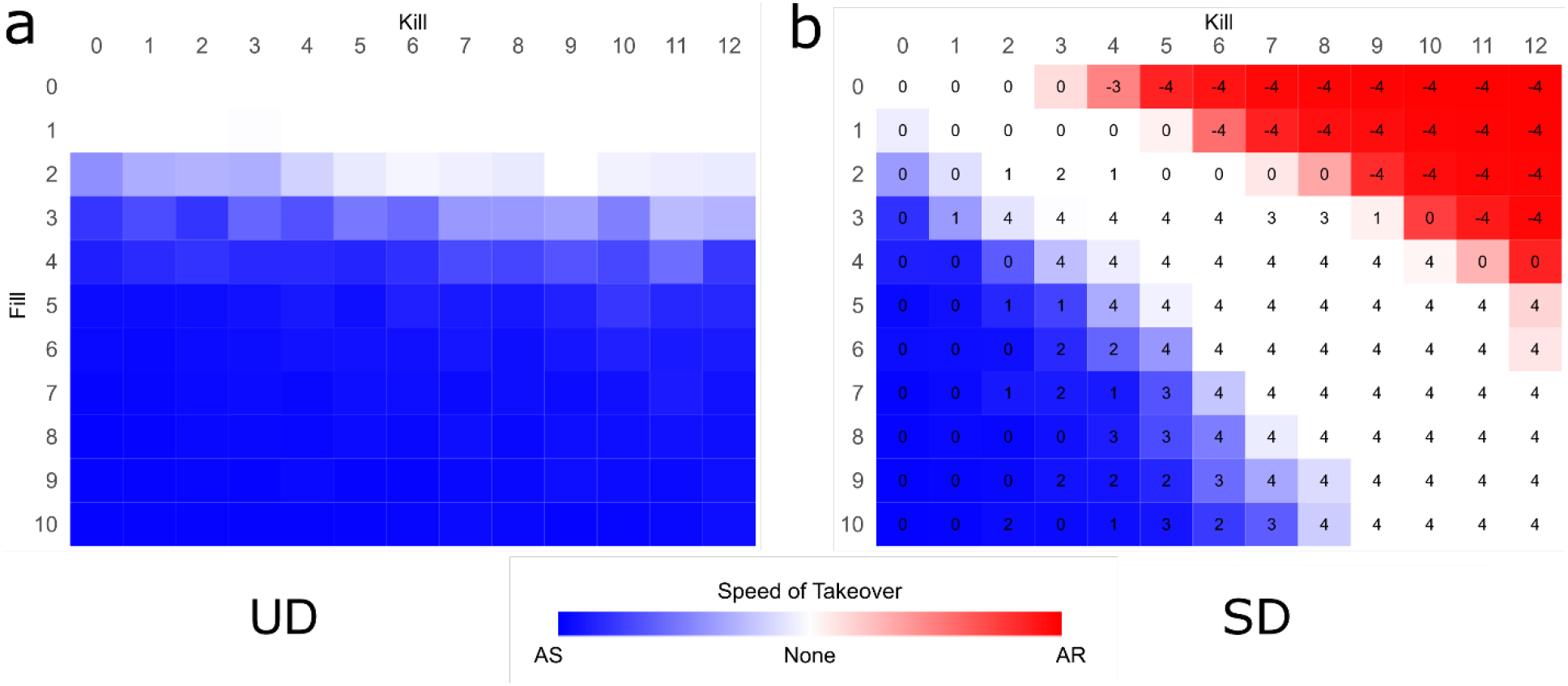
SD outperforms UD in mutant control. Heatmaps show speed and direction of mutant takeover for UD (A) and SD (B). Each tile reflects the mean takeover speed for 12 replicates (0-120,000 cycles), with red and blue colouring indicating takeover direction, and white tiles reflecting no mutant takeover over the timecourse. Increasing Fill values reflect additional growth advantage (probability of filling gaps) for cells with higher sensitivity, and increasing Kill values represent higher probability that cells with different sensitivity to their neighbours will be killed by them (except for slower cells in SD where higher Kill values reduce the chance of them being killed, see Figure 2C). Numbers reflect results of two-tailed T-tests comparing each Kill-Fill combination between SD and UD. Number 0 indicates SD was not significantly different to UD; 1 indicates p < 0.05; 2 indicates p < 0.01; 3 indicates p < 0.001; 4 indicates p < 0.0001. Numbers are negative if takeover occurred earlier than in UD. Speed of takeover reflects the number of cycles before a mutant cell type exceeds the number of wildtype.

As shown by the average sensitivity of cells over the time course (Supplementary Figure 1), reducing the rates of replication error and mutation (non-replication) can decelerate/prevent takeover within the time limit, while increased rates accelerate takeover, but neither alters the direction of takeover except at such high error levels where no cell can gain any advantage. This suggests that even though we have used high error rates to accelerate the models, the same outcomes will hold true for slower mutation rates in vivo over longer periods.

As UD is no less efficient at killing faster cells, the increased capacity of SD to prevent AS takeover must reflect the reduced capacity of the AS mutants to kill slower cells, which includes wildtype cells. Thus, the survival advantage of slower cells provides an important force of counterselection against the growth advantage of faster cells. However, as indicated by the low *Fill*, high *Kill* outcomes, if the counterselection was too strong for the range of viable mutations (*Fill* value), then SD could cause the spread of AR mutants. If SDT is correct, the result would be metabolic slowdown, declining tissue function, and thus ageing.

However, it is equally clear that by balancing the selective and counterselective forces, wildtype cells could retain their dominance with SD. While it is impossible to know whether running these simulations indefinitely would result in mutant takeover, organisms need only delay takeover until they are most likely to have died from extrinsic (or even other intrinsic) causes, and maintaining balance is possible at least for finite periods.

### Selective Destruction as an Inducer of Ageing

Initially this appears a strong argument against SDT. If selection and counterselection can be balanced, then SD need not cause the spread of AR mutants to prevent the spread of AS mutants. Negligible senescence could therefore be a viable life history trait and ageing an unnecessary one. However, simply preventing AS cell dominance and takeover may not be sufficient to remove the threat these mutants pose. SDT predicts that AS mutants induce death principally through CFOA. Under conditions where AS mutants arise and are cleared, the risk associated with them will depend on their numerousness at any given time (more AS cells at time T, will result in higher risk at time T, even if their ultimate clearance has removed this risk by time T + 1). Therefore, we firstly assessed the total abundance of AS cells over the timecourse for conditions associated with WT dominance (and negligible senescence) compared with those that allow the spread of AR mutants (and ageing). As shown in Figure 5A, the ageing life history trait is associated with significantly fewer AS mutants over the timecourse (as determined by T-test between highest negligible senescence *Kill* value and lowest ageing *Kill* value), and thus reduced risk of CFOA. We further considered additional relevant factors. Using functions described in the Methods, we created a tumour risk value (TRV) and fibrosis risk value (FRV). As overactivity is cell-type specific, we did not include it here.

**Figure 5.**
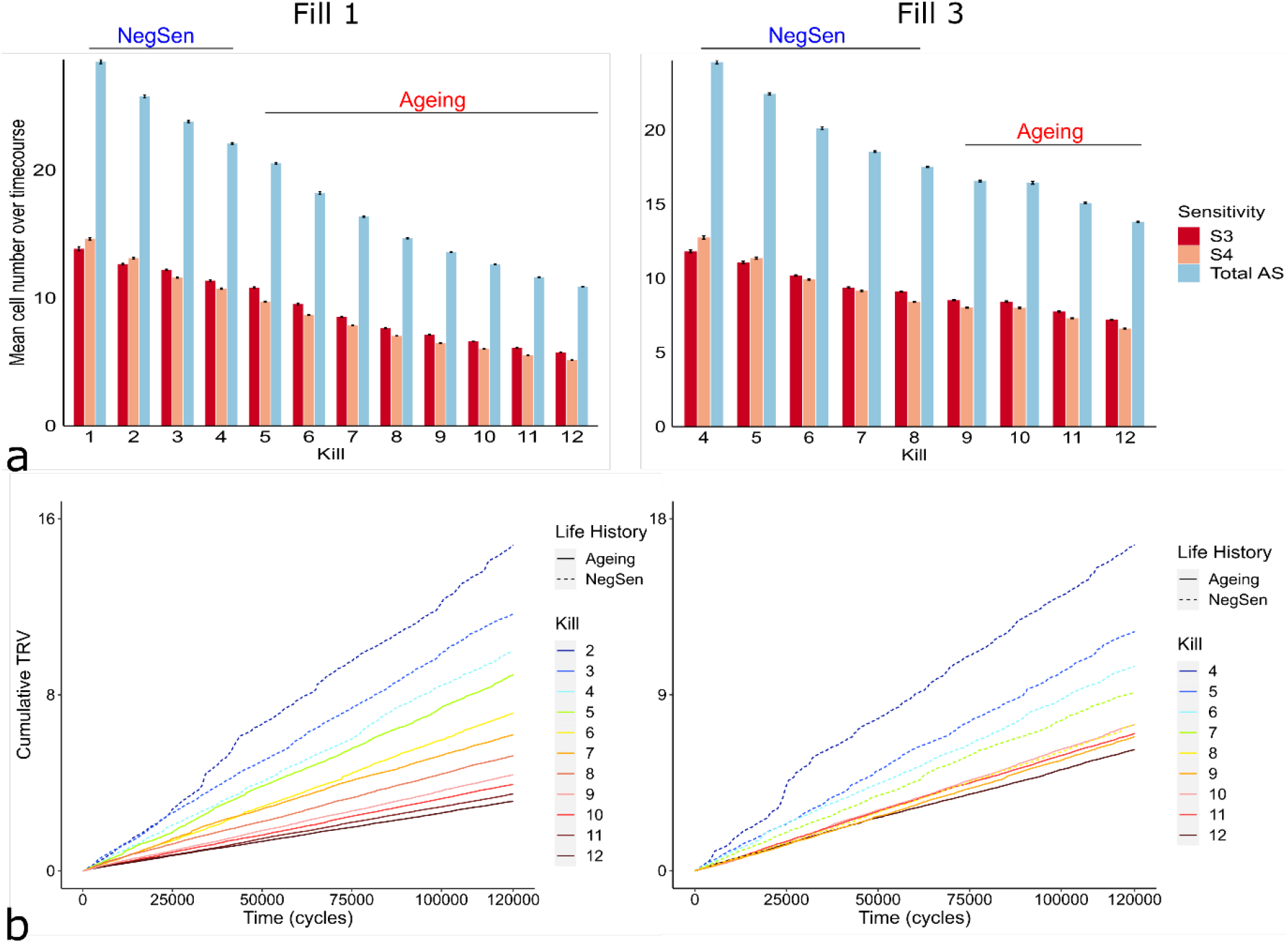
Ageing provides increased protection from AS mutants. (A) Mean number of each sensitivity of AS mutant (and total), and (B) Cumulative TRV over 120,000 cycle timecourse for the range of Kill values corresponding to negligible senescence and ageing life history traits (that result from wildtype and AR dominance, respectively). Two Fill values are shown reflecting different levels of growth advantage for AS mutants. NegSen, negligible senescence; S3, cells with sensitivity value (GM) 3; S4, cells with sensitivity value (GM) 4; Total AS, combined S3 and S4; TRV, tumour risk value (see Methods).

For TRV, we considered that the longer a specific mutant cell persisted and the more clones it produced, the more likely it would be to receive the series of additional mutations required for transformation. As shown in Figure 5B, the cumulative TRV is significantly greater for *Kill* values producing WT dominance than for *Kill* values producing AR dominance, and regression analysis of the highest *Kill* value for negligibly senescing populations and lowest *Kill* value for ageing populations showed the additional risk was significant (p < 0.001 for *Fill* 1 and 3). Thus, negligibly senescing organisms may be associated with greater risk of cancer than ageing ones. For FRV, we considered that the main determinant of fibrosis was the clustering of AS mutants, as wounding/inflammation at those clusters would be more likely to trigger cascades that led to self-sustaining populations of fibrotic cells (Adler et al. 2020). Risk at each cycle was therefore calculated as the sum of the sizes of the mutant clusters squared. Equally, the cumulative FRV rose more steeply for *Kill* values producing WT dominance than AR dominance (Supplementary Figure 2). The results clearly demonstrate that increasing counterselection beyond wildtype dominance into AR dominance provides further protection from CFOA.

### Positive Selection of Ageing by Selective Destruction

We have already described how the spread of AS mutants results in serious fitness costs in the form of CFOA; however, if their control by SD allows AR mutants to spread, then this will also cause a shift in tissue homeostasis, which might result in its own fitness costs. Here we are proposing that it results in ageing, which would therefore reflect a shift toward slower metabolism as cells become increasingly resistant to growth and replication signals over time. Although the AS and AR mutations themselves are somatic, the system of control will be governed by genes which can be inherited.

Alleles/genes which evolve to increase counterselection by strengthening SD will undergo positive selection if the fitness benefit they provide in the form of reduced risk of CFOA is greater than the fitness cost of ageing. To do this, we separated the impacts of ageing and CFOA into extrinsic and intrinsic risks as defined in the Methods. The fitness benefits from SD would thus reflect a reduction in intrinsic risk (as CFOA will induce death largely independent of extrinsic factors), while ageing first reduces organisms’ capacity to respond to extrinsic threats (such as disease and predators) and then later increases intrinsic risk in the form of age-related disease. We therefore constructed a gene selection model of organisms with three genes, each with two alleles that either strengthened SD (S alleles) or relied on UD (U alleles). Although this is likely an oversimplification, we considered it sufficient to illustrate the impacts of mutant control on intrinsic and extrinsic fitness that work beneath the additional variables complicating these relationships in actual in vivo conditions. As described in our tissue model (Figure 4), a purely UD control mechanism is insufficient to prevent AS dominance/spread and death from CFOA. Therefore, in our gene selection model, individuals with UUU genotype had increasing intrinsic risk from birth, as shown in Figure 3. A single S allele provided SD strong enough to prevent AS spread, but not sufficiently strong to induce AR spread. This represented the balanced condition that allowed wildtype dominance throughout, which could result in negligible senescence. As indicated in Figure 5, these individuals would have a higher basal risk of CFOA than if SD was stronger, but the clearance of AS mutants would prevent risk from increasing, as with UUU individuals. Individuals with two or three S alleles would have strengthened SD sufficiently to allow the spread of AR mutants, and thus ageing would result. We modelled this as increasing extrinsic risk commencing after an initial growth period at an age determined by the variable, *Peak*. Later, intrinsic risk also increased after the age determined by the variable, *Rapid decline* (reflecting age-related diseases). Individuals with SSS genotype reached this age earlier than those SSU genotype, and had accelerated increase in extrinsic risk after peak age. Thus, the model had four phenotypes reflecting four genotypes, which in turn reflected the strength of SD and the dominance of the AS/AR/wildtype cells that resulted, as shown in Figure 3. Further model details are found in the Methods.

For simplicity, all increases in risk were linear, reflecting the survival functions in Supplementary Table 1. We used a comprehensive range of intrinsic and extrinsic survivability values (see Methods) ranging from zero (certain death) to negligible risk (causing exponential population increase for extrinsic survivability). Risk associated with genotype is reduced as survivability increases, allowing us to see, in a spectrum shifting the relative importance of intrinsic and extrinsic factors, which points favour which life history traits (ageing and negligible senescence).

As shown in Figure 6A and B, when looking at the average abundance of individuals with each genotype over a 100,000 cycle time course (as percentage), the results showed conditions where negligible senescence, ageing, and rapid ageing phenotypes produced the greatest fitness. This was also true across a range of combinations of peak fitness age (*Peak*, where ageing began to affect fitness) and *Mating age* (where offspring could first be produced) tested across different intrinsic and extrinsic survivability values (Figure 6C-E), and importantly was unaffected by whether the population started with three U alleles (increasing CFOA, Figure 6A) or three S alleles (rapid ageing, Figure 6B), suggesting this was the result of positive selection rather than drift.

**Figure 6.**
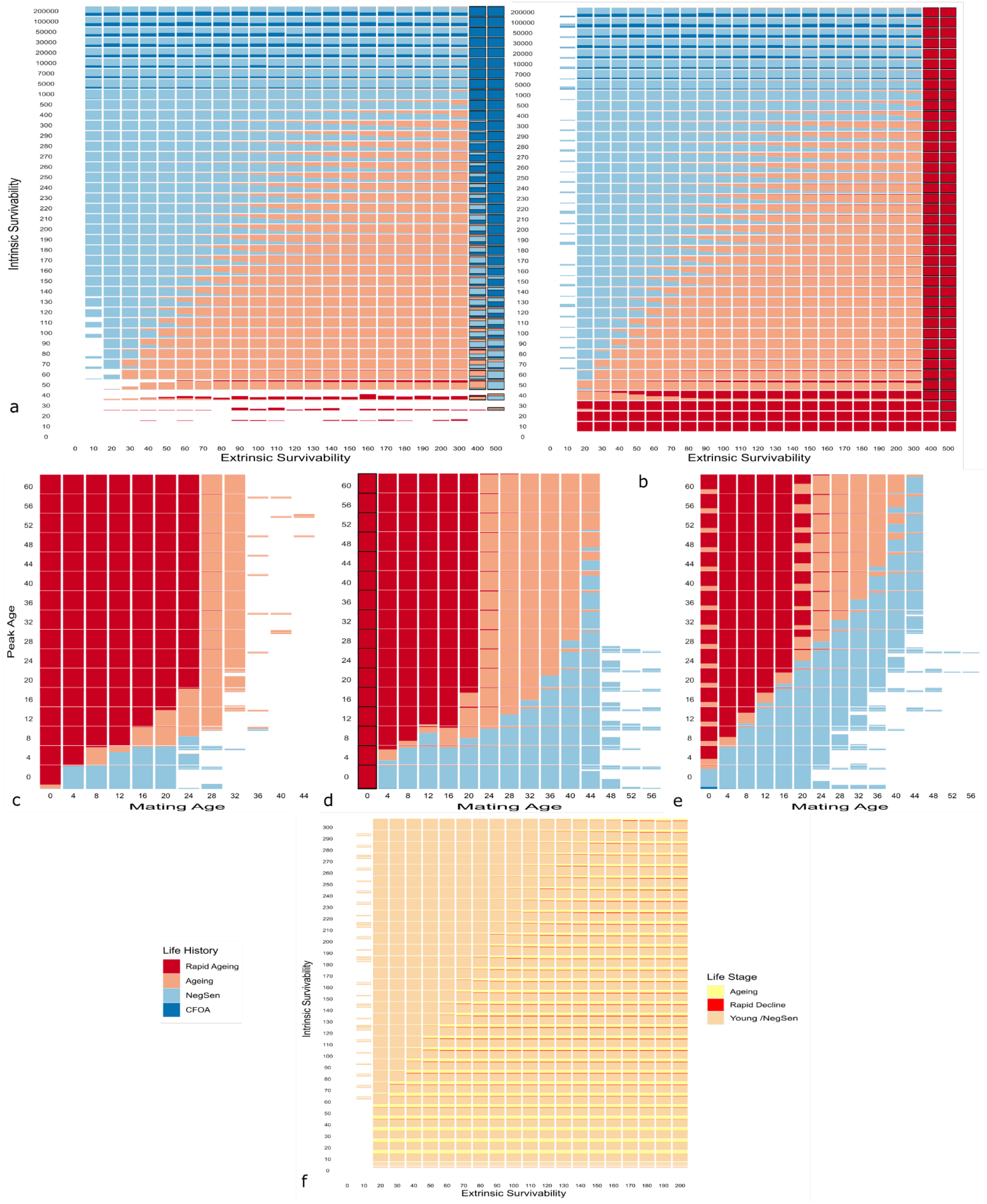
SD and ageing undergo positive selection under various conditions. (A and B) Each tile shows the mean abundance (as percentage) of agents with the four life history traits over the 100,000 cycle time course. The emergent life history traits reflect the allelic combinations shown in Figure 3 and Supplementary Table 1. The survivability ranges from zero to negligible risk for both intrinsic and extrinsic factors. A survivability value of Y means there is an X in Y chance of death at each cycle, where X is the risk value of an individual within the population (determined by its allelic combination). In negligibly senescing populations, X is determined by population size for extrinsic survivability, and number of S alleles for intrinsic survivability. In ageing populations, age beyond peak fitness and Rapid decline will also affect extrinsic and intrinsic X values, respectively. White space indicates populations went extinct, black borders indicate populations underwent exponential increase and simulations were ended at 10,000 agents. Units are all arbitrary units. Peak & Mating age, 10; Rapid decline, 20. (A) Starting populations are uniform UUU alleles (for UD). (B) Starting populations are uniform SSS alleles (for rapid ageing with strong SD). (C, D and E) Tiles represent mean abundance (as percentage) of agents for each life history trait over the time course for different peak ages (after which ageing begins to affect fitness) and Mating age, before which individuals cannot produce offspring. Rapid decline, 50. (C) Intrinsic survivability 50 and extrinsic survivability 50 (close to rapid ageing dominance in A and B). (D) Intrinsic survivability 150 and extrinsic survivability 150 (ageing dominance). (E) Intrinsic survivability 250 and extrinsic survivability 50 (negligible senescence dominance). (F) Mean abundance of agents in peak fitness (both pre-ageing and negligibly senescing), ageing (decreasing extrinsic fitness), and rapid decline (decreasing intrinsic and extrinsic fitness) across the same range of extrinsic and intrinsic mortality values in A and B. NegSen, negligible senescence.

Thus, ageing via SD could be expected to undergo positive selection and spread in multiple environments, particularly as intrinsic risk (from CFOA) increases. Further, we looked at the abundance of agents at different life stages; either young (pre-ageing or agents with negligible senescence), ageing (Age > *Peak*, ie. past peak fitness with increasing extrinsic risk), and in rapid decline (Age > *Rapid Decline*, i.e. suffering increased intrinsic risk from age-related diseases). Figure 6F shows that in most scenarios where ageing was the dominant life history trait, a fraction of the population had passed peak fitness and were undergoing decline, with some individuals even entering the period of rapid decline. Thus, ageing by SD could be expected to undergo positive selection even in populations where individuals were reaching ages where fitness began to decline, consistent with observations that ageing occurs in the wild (Nussey et al. 2013).

Importantly, this toy model leaves out many details affecting intrinsic and extrinsic mortality such as accumulating mutations, which will increase cancer risk independent of these mutant clearance mechanisms (Hanson et al. 2015), and the initially high extrinsic risk for younger organisms (Kinzina et al. 2019). The results have obvious limitations when extrapolating outcomes for real life populations; however, they demonstrate clearly that if negligible senescence is accompanied by a higher basal risk from intrinsic factors, it could cause the selection of genes which induce ageing (and its associated increasing extrinsic and intrinsic risks), implicating that ageing and the rate of ageing could depend on the strength of SD. Changes in the number of genes or alleles and the proportion to which they strengthen or weaken SD should not shift this fundamental relationship, but additional factors will complicate it, shifting the potential of SD to affect fitness. Thus, while these results are sufficient to demonstrate the plausibility of SDT, further work is required to evidence the occurrence of SD in real-world populations.

### Selective Destruction and Molecular Damage

It might be argued that because the generation of AS and AR mutants involves molecular damage that such damage could be prevented by the upregulation of maintenance, in line with DST. However, the level of damage required to induce a change in growth sensitivity could be as simple as a single point mutation – multiple different point mutations in RAS are associated with tumorigenesis (Miller and Miller 2012). Thus, SD does not require damage *accumulation*, and predicts that upregulating maintenance will only extend lifespan to the extent that it can significantly slow single base mutations in the DNA. Plausibly, upregulating maintenance mechanisms to such an extent would delay the cell cycle and transcriptional machinery to a degree that would reduce fitness even aside from the associated energetic costs. Therefore, while molecular damage likely plays a significant role in selective destruction, the energetic costs of maintenance need not. The fundamental cause would thus be different to damage-accumulation-based theories such as DST.

In addition, while we have been referring to changes in growth signalling as mutations, they are likely to reflect both genetic and epigenetic changes. Epigenetic changes are heritable and semi-permanent, so could provide long lasting changes in growth signalling. In our previous models we have assumed that SD works primarily by killing the surrounding cells. However, this mechanism is obviously inefficient. We therefore considered a scenario where a certain percentage of neighbours identifying more growth sensitive cells would induce epigenetic changes to slow them down (rather than killing them). Slower cells would still be removed as before.

As shown in Figure 7, 50% and 100% conversion of faster cells to one or two sensitivity levels below their current level were significantly and increasingly more effective at controlling AS mutants than lower levels of conversion, as determined by two tailed T-tests. As such, the spread of AR mutants may not reflect mutation or molecular damage at all, but an evolved response to damage that could occur even in the absence of it. This finding has particular significance because programmatic theories of ageing are widely deemed to be evolutionarily implausible (Kirkwood 2005; Kirkwood 1977), unlikely to undergo the kind of positive selection that would make ageing a universal phenomenon. However, if cells with lower growth sensitivity are slowing their faster growing neighbouring cells over time, we must conclude that an epigenetic program of metabolic slowdown could evolve, undergo positive selection, and induce ageing. The rate of ageing would be determined both by the strength of the SD and the rate of conversion. The only remaining question is whether this is actually happening.

**Figure 7.**
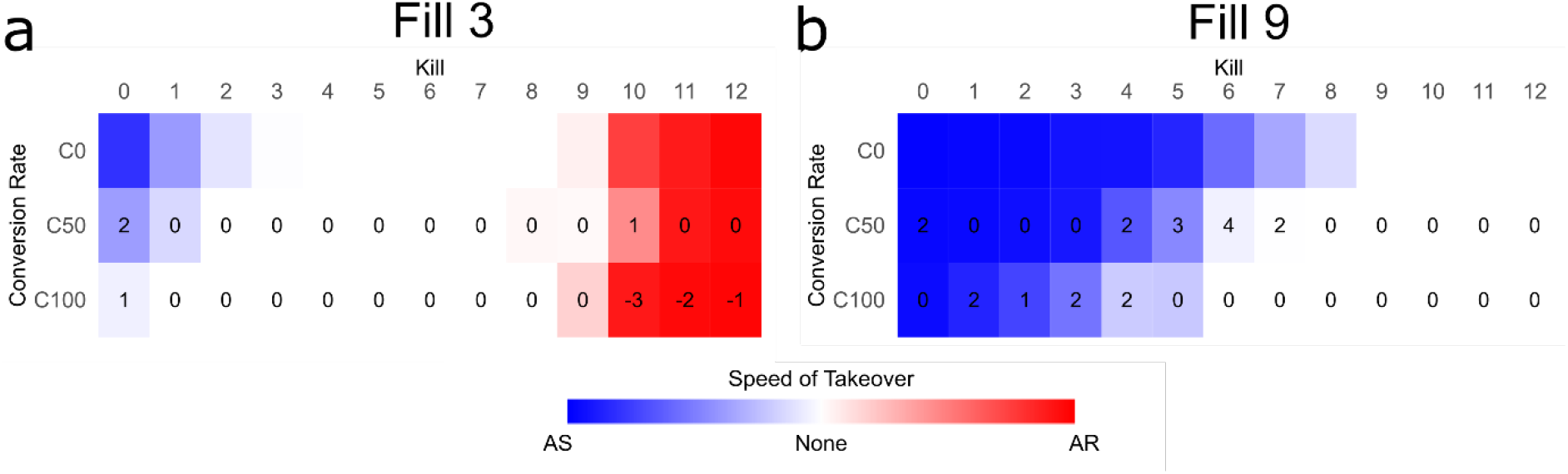
Epigenetic retardation of cells outperforms cell killing at AS mutant control. Heatmaps show speed and direction of mutant takeover for SD with different rates of conversion to slower sensitivity. Speed of takeover reflects the average value for 12 replicates. Numbers reflect results of two-tailed T-tests comparing each Kill-Fill combination between the conditions. For T-tests, SD with 50% conversion (and 50% killing, SD50) was compared to SD with zero conversion (100% killing, SD0), and SD with 100% conversion was compared to SD50. Number 0 indicates no significant change; 1 indicates p< 0.05; 2 indicates p < 0.01; 3 indicates p < 0.001; 4 indicates p < 0.0001. Numbers are negative if takeover occurred earlier with higher percentage conversion. (A) is for Fill value 3, (B) for Fill value 9, showing results for AS mutant cells with relatively low and high growth advantage, respectively.

## Discussion

Here we demonstrate the advantage of SD and its implications. Although the theoretical nature of this work necessitates that the models may not reflect how SD works in vivo, the simplicity of our system demonstrates clearly that if mutants are arising with even the weakest growth advantage, a counterselective force, such as SD, is necessary to control them. Indeed, it was the absence of such a control mechanism that led Nelson and Masel (2017) to conclude that ageing was an inevitable result of multicellularity: their model showed that non-cooperative mutant cells would inevitably overcome the wildtype cells *because it included no counterselective force*, consistent with our results for UD (Figure 4A). Our combined results underline the critical nature of the evolution of SD for multicellular life. Whether it works via Notch as we suggest, or immune regulation, or another mechanism is still hypothetical, but even its existence in some form has large implications. If strengthening this force until it results in the spread of slower mutants provides continuing advantage, as we suggest it might, then this will result in a shift in homeostasis over time. To our knowledge this is the first mechanistic assessment of why such a process might evolve and thus contribute to ageing, and this initial cause would be fundamentally different to the costs of maintenance (DST). While we do not doubt that accumulating damage plays an important role in ageing, the widespread failure of anti-ageing treatments aimed at removing molecular damage may implicate additional mechanisms. If ageing is, even in part, a metabolic slowdown from SD, then even if this causes molecular damage to accumulate at late stages, removing this damage will have short term effects at best, as it is the altered functionality of cells which causes them, for example, to produce misfolded proteins and not remove them. Thus, some accumulated damage may not reflect the slow build-up over time, but a shift in homeostasis from clearance to accumulation, suggesting it will quickly return to previous levels when treatments are ceased. In the case of alagebrium to treat arterial inflexibility by the removal of advanced glycation end products (AGEs), De Grey (2007) describes the reason “treated animals become inflexible again so quickly after withdrawal of the drug “ as possibly due to exposure of a highly reactive carbonyl group within the degraded AGE products, but it could equally reflect that older individuals have a higher equilibrium level of AGEs, and ceasing treatment naturally and swiftly causes return to this pre-treatment point. Conversely, DST predicts that activities such as smoking which increase molecular damage would be expected to affect mortality long after they cease, due to the accumulated damage they cause, but the opposite is true (Alberg et al. 2013). Importantly, smoking increases clonal expansion of cells with driver mutations (causing cancer), which are cleared once individuals cease smoking (Yoshida et al. 2020), as we have shown for AS cells in the models of SD.

Unlike treatments designed to remove accumulated molecular damage, which might be expected to have little downside, treatments that slow ageing by weakening SD could be expected to increase the risk of early death by CFOA. We therefore predicted that there would be some longevity treatments that induced a degree of early deaths, and sure enough multiple calorie restriction (CR) studies in mice (Cameron et al. 2012; Koopman et al. 2016; Mitchell et al. 2016; Yu et al. 2019) showed Kaplan-Meier curves with early dips in survival in the CR groups compared with control, as did the Kaplan-Meier curves presented by Mattison et al. (2017) for the two main primate studies. This occurred even though the survivors of CR went on to live longer and/or healthier lives than controls. Therefore, a key impact of SDT is that curing ageing need not produce immortality. Instead, negligible senescence may simply reflect a higher basal risk of death. A recent review by Frenk and Houseley (2018) examined a collection of meta-analyses and systematic reviews of gene expression changes with age, identifying six hallmarks reflecting consistent changes across multiple species including worms, flies, mice, and humans, as well as different organs within these species. The first and second most common hallmarks were the downregulation of mitochondrial genes and the downregulation of protein synthesis machinery, particularly ribosomal proteins and biogenesis factors, which is highly consistent with a metabolic slowdown resulting from SD. Another hallmark was the reduction in growth factor signalling. The reviewed studies demonstrated reduced expression of genes associated with cell growth in human and worm muscle (Ma et al. 2016; Zahn et al. 2006); reduced expression of IGF-1 and GH pathway genes in mouse liver (Schumacher et al. 2008); and DNA methylation of cell cycle genes in β cells (Avrahami et al. 2015). All these changes are highly consistent with SDT. Notably, rates of cancer generally increase with age which may appear inconsistent with a mechanism of ageing designed to remove potentially tumorigenic cells. However, we are not suggesting that even the strongest SD could remove all mutant cells. As such, successive insults to the genome would still cause increasing cancer rates with age, but at a much slower rate than would be the case without SD. Interestingly, studies suggest that the rate of cell division in humans declines with age (Tomasetti et al. 2019), which has been proposed to explain the reduced rates of cancer in the oldest old (Hanson et al.2015).

Perhaps the main challenge for SDT is that in mainly post-mitotic organisms such as C. elegans and Drosophila, a mechanism of ageing that relies mainly on cell proliferation is much less likely to play a role. However, the opposite is also plausible: these organisms have evolved to become post-mitotic to escape the dangers of AS mutants, making the post-mitotic life history trait an extreme form of SD which essentially removes every proliferating cell. Although obviously speculative, it is difficult to imagine why the post-mitotic life history trait would be beneficial except by preventing the dangers associated with cell division (i.e. CFOA). Consistently, while adult Drosophila have cancers limited to their gut (the only remaining mitotic tissue), the mitotic larval Drosophila can get cancer in multiple tissues (Eichenlaub et al. 2016; Suijkerbuijk et al. 2016). This strategy of SD via postmitosis would totally prevent mutant spread at the cost of the organisms’ ability both to regulate outputs through organ size (Karin et al. 2016) and to remove damaged cells and replace them with new ones. Post-mitosis would therefore potentially induce mechanisms of ageing which were avoided in their single-celled ancestors like yeast, which essentially divide away the damage (although this is conventionally referred to as ageing) and need not occur in organisms that maintained or re-evolved mitotic somas. As modelled by Nelson and Masel (2017), damage accumulation would reflect the lack of cell competition (and removal) that would allow the low-vigor/damaged cells to persist in post-mitotic organisms, driving a different form of ageing to systems which retained cell competition. If SDT is correct, we would predict that molecular damage accumulation may prove less relevant to ageing in mitotic than post-mitotic organisms.

We believe that SDT provides the first theory which could explain both the evolution and mechanism of ageing which would also undergo positive selection independent of molecular damage accumulation and the associated energetic costs. The necessity of studies to frame their results within the sphere of accumulating damage has created a myopic view of ageing, so a second hypothesis could be a huge boon to the field. That said, we accept that many of the concepts put forward here are in their nascent stages. The idea of neighbouring cells controlling each other’s growth rate is currently hypothetical. We have suggested Notch signalling may be a key part of this process and shown that mutations in this network are clearly linked with loss of mutant control and cancer, but there is still considerable experimental work to be done here, and the relevance of SD to ageing will come down in large part to the results of these experiments. While mutant control must exist in some form to prevent the result indicated by Nelson and Masel (2017), it is possible that AS mutants requiring non-autonomous control do not occur in sufficient quantities to merit a system of SD. The rates of mutation affecting growth sensitivity used in our simulations were artificially high for the sake of expediency.

Notably, both mutations and epimutations could generate AS mutants, and the latter may be orders of magnitude more frequent than the former (van der Graaf et al. 2015). Indeed, the very idea of SD through epigenetic conversion may require redefinition of “epimutation “. Initially defined by Holliday as “aberrant patterns of DNA methylation that cause the silencing of a normally expressed gene, “ or the contrary aberrant “ectopic expression “ (Holliday et al. 1990), within the context of SD this should at least be subdivided into those that are adverse reflecting chance conditions, and those that are deliberately induced to slow the metabolism of potentially dangerous cells (or otherwise spread by the force of this counterselection). If SD is correct, the latter kind likely produce the epigenetic clocks that predict both chronological and biological age, while the former will represent the increasing epigenetic noise.

Evidence that clonal expansion occurs with age and contributes to cancer is continually increasing, recently reviewed by Kakiuchi and Ogawa (2021), who described the spread of clonal mutants across multiple tissues, including one study where “virtually all endometrial glands were replaced by one or more driver-mutated clones “ in women by the age of 50 (Moore et al. 2020).

To conclude, we have shown that SD provides a more effective mutant control mechanism than UD, and described how reducing the risk of cancer, fibrosis, and overactivity (CFOA) might explain why organisms strengthen SD to the point that it begins to allow the spread of AR mutants, with epigenetic growth suppression potentially also contributing to the metabolic slowdown. SD could therefore provide the proximal cause of ageing.

## Acknowledgements

We would like to acknowledge Professor Tom Kirkwood and Dr Viktor Korolchuk for developmental input. Images were created using BioRender.com. This work was supported by the Novo Nordisk Fonden Challenge Programme: Harnessing the Power of Big Data to Address the Societal Challenge of Aging (grant number NNF17OC0027812).

## Author contributions

Author contributions: James Wordsworth developed the theory and models, and wrote the manuscript with input and supervision from Daryl Shanley. Hannah O’Keefe contributed to NetLogo model development and Peter Clark helped with cluster work.

