## Supplementary Fig1 and 2 for "The damage-independent evolution of ageing by selective destruction"

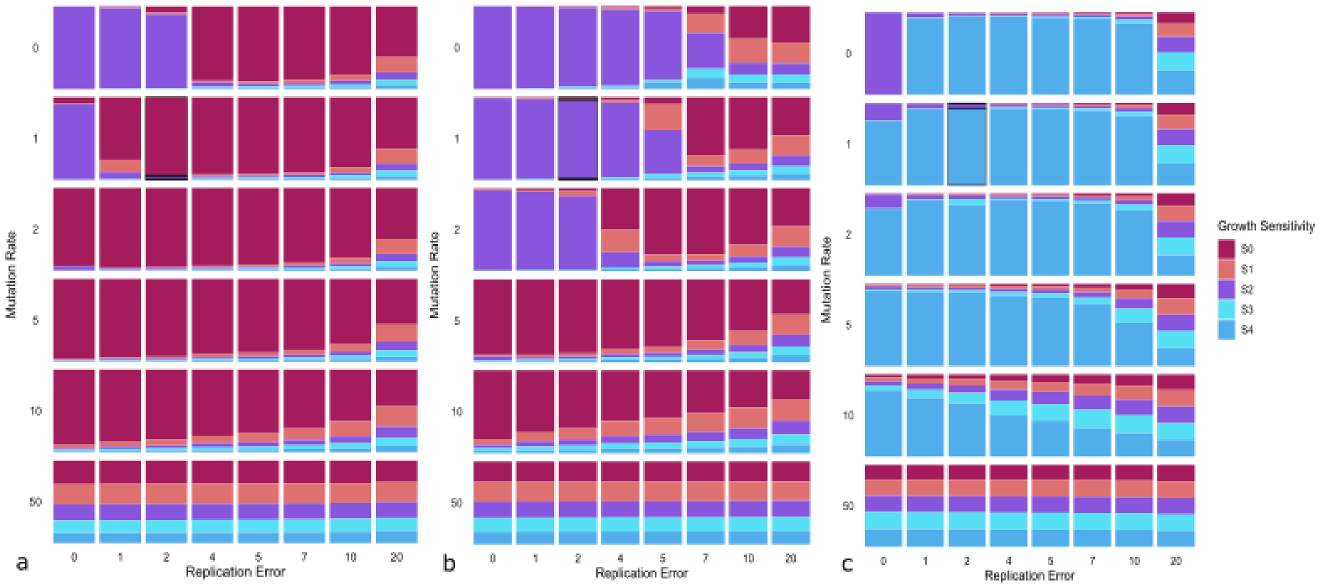


*Figure S1|Effect of varying replication error and mutation rate on the percentage of cells with different growth sensitivities (S1-S4). Each tile reflects the mean of 12 replicates for the abundance of cells with different growth sensitivities (as percentage) over the 120,000 cycle timecourse. Three different Kill and Fill combinations in the SD model were used corresponding to: (A) AR mutant takeover (Fill 2 Kill 11); (B) Wildtype dominance (Fill 5 Kill 8); and (C) AS mutant takeover (Fill 8 Kill2). Mutation rate 1% and replication error rate 2% were used in other simulations.*

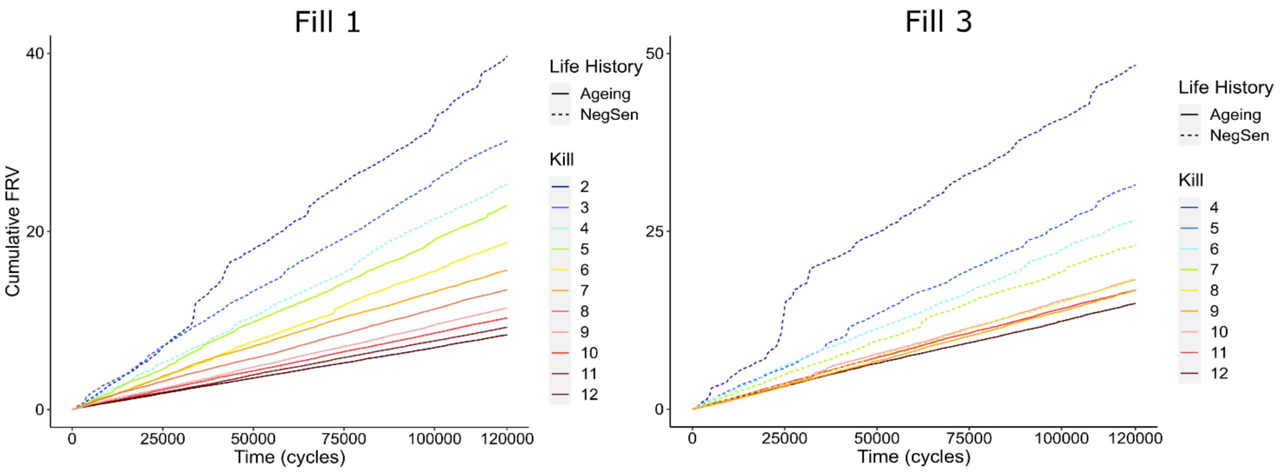


*Figure S2 | Cumulative FRV over 120,000 cycle timecourse for the range of Kill values corresponding to negligible senescence and ageing life history traits (that result from wildtype and AR dominance, respectively). Two Fill values are shown reflecting different levels of growth advantage for AS mutants. FRV, fibrosis risk value (see Methods)NegSen, negligible senescence; S3, cells with sensitivity value 3; S4, cells with sensitivity value 4; Total AS, combined S3 and S4. Regression analysis of the highest Kill value for negligibly senescing populations and lowest Kill value for ageing populations showed a significant increase in fibrosis risk for negligibly senescing populations (p < 0.001 for Fill 1 and 3).*

| Alleles | AgeTot (AT) | AgeRate (AgR) | Dominance | Emergent life history trait | Survival Functions | Function summary |
|---|---|---|---|---|---|---|
| UUU | 3 | 0 | AS mutant | Aberrant growth diseases from increasing CFOA risk | Die if X' < 1 | Constant extrinsic risk |
| | | | | | Die if Y' < AT + Age | Intrinsic risk increases with age from birth |
| SUU | 2 | 0 | Wildtype | Negligible senescence | Die if X' < 1 | Constant extrinsic risk |
| | | | | | Die if Y' < AT | Constant intrinsic risk |
| SSU | 1 | 1 | AR mutant | Ageing | if Age >= Peak: $X_{(t+1)}$ = ($X_t$ - AgR)<br>Die if X' < 1 | Extrinsic risk begins increasing after peak age |
| | | | | | **if** Age < RD / AR: Die if Y' < AT<br>**else** die if Y' < AT + (Age - RD / AgR) | Intrinsic risk begins increasing at older age (RD) |
| SSS | 0 | 2 | AR mutant | Rapid ageing | if Age >= Peak: $X_{(t+1)}$ = $X_t$ – AgR<br>Die if X' < 1 | Extrinsic risk begins steeply increasing after peak age |
| | | | | | **if** Age < RD / AR: Die if Y' < AT<br>**else** die if Y' < AT + (Age - RD / AgR) | Intrinsic risk begins increasing at older age (RD/2) |

*Table S1 | Survival functions for the different genotypes of the gene selection model. Each cycle, each individual would 'Die if' they met the criteria corresponding to their genotype and age. X is the value of extrinsic survivability as determined by the model, and Y is the value of intrinsic survivability. Both are stochastic reflecting chance conditions and occurrences, thus **X'** and **Y'** refer to a randomly selected number between 0 and X or Y, so as X and Y increase, risk of death (Die if) decreases. The Peak variable determines the age after which extrinsic fitness begins to decline. RD, rapid decline is the variable determining the age at which ageing begins to have significant effect directly on intrinsic mortality. AgeTot (AT) is the number of U alleles, and AgeRate (AgR) is the rate of ageing determined by the number of S alleles. AS, aberrently sensitive to growth; AR, aberrantly resistant to growth; CFOA, cancer fibrosis and overactivity.*
